# Alpha-synuclein modulates the positioning of endolysosomes in melanoma cells

**DOI:** 10.1101/2025.03.22.644754

**Authors:** Nirjhar M. Aloy, Maria Ericsson, Brandon Hartman, Stephan N. Witt

## Abstract

The Parkinson’s disease-associated protein, alpha-synuclein (α-syn; *SNCA*) is suspected of promoting melanoma progression. We recently knocked out *SNCA* in the human cutaneous melanoma cell line SK-MEL-28 to try to deduce the role of α-syn in melanoma progression. Compared to control cells, the SK-MEL-28 *SNCA*-knockout (KO) cells have significantly inhibited growth, invasion, and migration, and the levels of the neural adhesion protein L1CAM and the transferrin receptor (TFR1) are significantly reduced. In this study, using transmission electron microscopy and immunofluorescence we show that SK-MEL-28 *SNCA*-KO cells relative to control cells exhibit an increased density of endolysosomes; increased perinuclear positioning of large (> 800 nm) endolysosomes; and decreased levels of the tetraspanins CD9 and CD81. Based on these results, we infer that α-syn disrupts the balance between anterograde and retrograde traffic; thus, we propose that α-syn is an accessory factor that that positively modulates the anterograde transport of endolysosomes and that loss of α-syn expression results events (i)-(iii). We infer that low levels of L1CAM and CD81 (and other membrane proteins) are likely the underlying reason for the significantly reduced invasiveness and migratory properties of SK-MEL-28 *SNCA*-KO cells.

## Introduction

There is a well-documented co-occurrence of Parkinson’s disease (PD) and melanoma. Such a co-occurrence seems unusual because PD is a neurodegenerative movement disorder that occurs from the progressive degeneration of midbrain dopaminergic neurons (1); whereas melanoma is a deadly cancer of the skin that arises from mutations in the pigment producing cells called melanocytes (2). Individuals with PD have a 75 % higher risk of melanoma (3), and, reciprocally, individuals with melanoma have a 50-85% higher risk of developing PD (4). Several genes may be involved in the co-occurrence of these two diseases (5–7) and one of them is *SNCA*, which is the gene that codes for the neuronal protein alpha-synuclein (α-syn) (8).

Melanocytes and neurons are both derived from the neural crest during development. Melanocytes and dopaminergic (DA) neurons have curious similarities (5), in that, each cell type (i) synthesizes pigment, i.e., melanocytes synthesize melanin and DA neurons synthesize neuromelanin; (ii) expresses tyrosine hydroxylase and tyrosinase, which are enzymes that catalyze the hydroxylation of tyrosine, the first step in the production of dopamine and/or pigment; and (iii) expresses the protein α-syn. This protein, α-syn, has a Dr. Jekyll and Mr. Hyde aspect, in that, α-syn appears to be involved in melanoma pathogenesis and progression is hence is pro-survival (9, 10), whereas α-syn, when aggregated (11) or at elevated concentrations due to gene triplications (12), causes neurodegeneration and hence is a cytotoxic protein.

In neurons, α-syn binds to presynaptic terminals (13) where it promotes synaptic vesicle clustering and SNARE complex assembly (14, 15). Other documented activities of α-syn include avid binding to membranes (16), lipids (17, 18), and DNA (19, 20), and promotion of endocytosis (21) and exocytosis (22, 23). In melanoma cells, the function of α-syn is less clear because this has not been an extensively studied area. We know that α-syn is expressed in aggressive melanomas (24), perhaps because of its pro-survival-like functions of modulating autophagy (25), vesicular trafficking (10, 26), and immune evasion (27, 28). α-syn also inhibits melanin synthesis, which makes melanocytes susceptible to damage from UV radiation (29). The use of a cutaneous melanoma cell line (SK-MEL-28), in which CRISP/Cas9 was used to knock out *SNCA*, has given new insights into the function of α-syn (10). Loss of α-syn expression in this melanoma cell line causes inhibition of growth, invasion, and migration and concomitantly decreases in the levels of cell surface proteins such as the transferrin receptor (TFR1), L1CAM, N-cadherin, and the tetraspanin CD81 (10, 26, 27), although the mechanistic details for these changes are unknown.

In this work, we present ultrastructural and immunofluorescence evidence that the loss of α-syn expression in *SNCA*-KO cells increases the density of endolysosomes, promotes the perinuclear clustering of large endolysosomes, and that the levels of proteins destined for the plasma membrane, CD9 and CD81, are reduced, whereas proteins, CD63 and Lamp1, which are embedded on the limiting membrane of the endolysosomes, build up.

## Results

### Endolysosomes accumulate in *SNCA*-KO melanoma cells

We previously reported increased lysosomal degradation of the cell-surface proteins TFR1 (10) and L1CAM (26) in SK-MEL-28 *SNCA*-KO melanoma cells compared to control cells. These findings prompted us to hypothesize that α-syn has a role in promoting the intracellular trafficking of these two cell-surface proteins, and likely others, through the endolysosomal system. To this end, we performed transmission electron microscopy (TEM) to observe the ultrastructure of control and *SNCA*-KO melanoma cells. Our objective was to determine whether α-syn expression (or lack of) affects the number, size, and positioning of endolysosomes. To simplify our analysis, we defined both multivesicular endosomes (MVEs) and endolysosomes commonly as endolysosomes because they frequently appear in a hybrid state (multivesicular and multilamellar) in the TEM images. There were previous reports describing MVEs as endolysosomes (30). We were particularly interested in the endolysosomes because previous reports had shown that the acid-hydrolase mediated degradation of proteins and other molecules take place in the endolysosomes and not in terminal lysosomes (31).

TEM was used to analyze the ultrastructure of the melanoma cells that were subjected to two different protocols. First, for cells grown and fixed on thermanox coverslips, we observed an accumulation of MVEs and endolysosomes in *SNCA*-KO cells compared to the control cells (Fig. 1A). A blinded quantification of the number of endolysosomes in *SNCA*-KO and control cells revealed a significant increase (*P* = 0.0076) in the number of endolysosomes per unit area in *SNCA*-KO cells compared to the control cells (Fig. 1B). Second, in a slightly different experimental protocol where ultrathin sections from cell pellets were prepared after detaching them gently with EDTA (Supplementary Fig. S1A), we observed heterogeneous endolysosomal structures.

**Figure 1.**
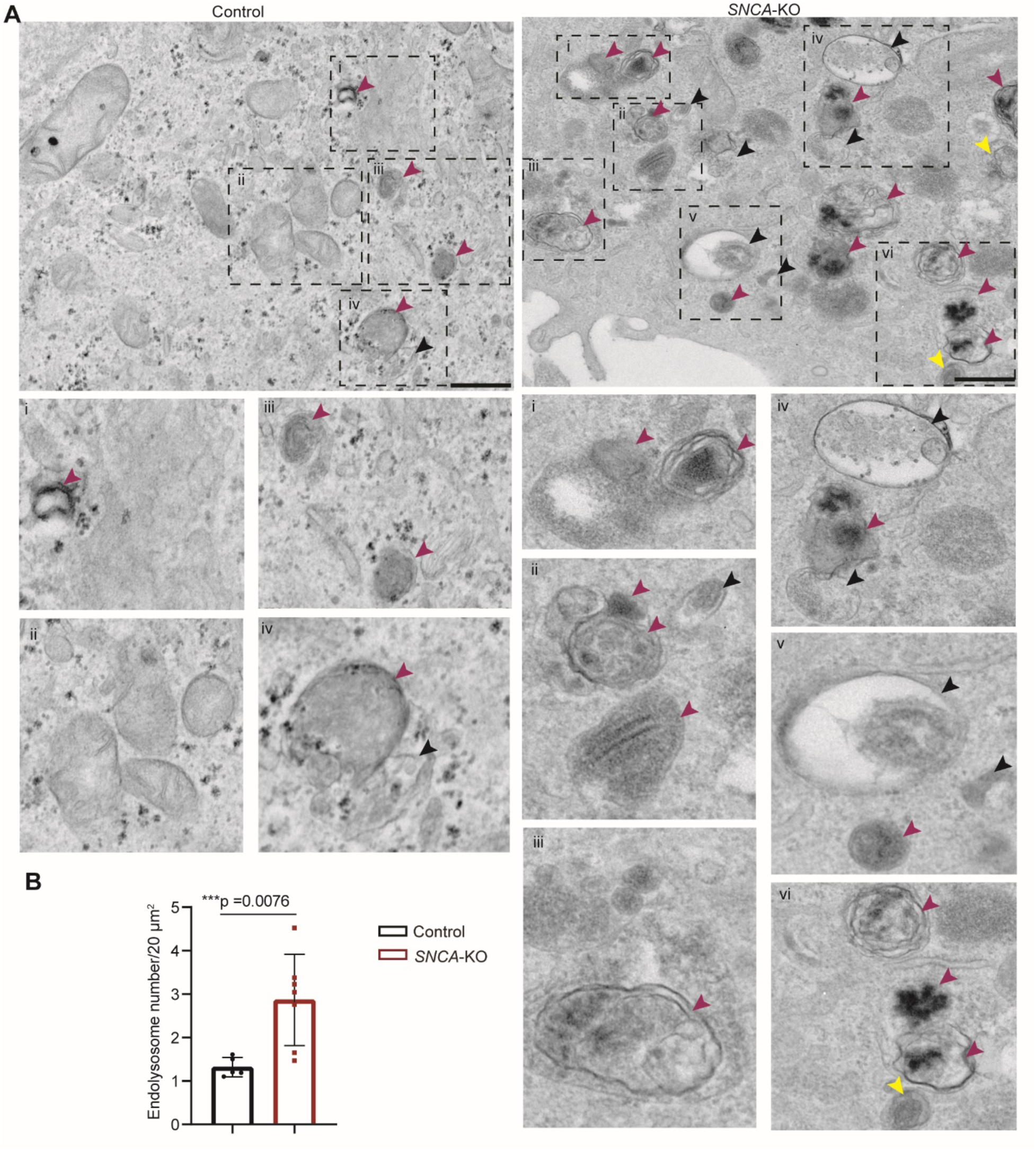
Endolysosomal structures are increased in number *SNCA*-KO cells. (A) Ultrastructural analysis by transmission electron microscopy of control and *SNCA*-KO cells showing the endolysosomal structures. Insets (numbered i-iv in the left panel and i-vi in the right panel) show the enlarged views (2x zoom using ImageJ) of the dashed rectangles with corresponding number from each cell type. Magenta arrows indicate endolysosomal structures (electron dense) and black arrows indicate MVEs. Representative images from two biological replicates are shown. Scale bars = 500 nm, Direct magnification 8000x. (B) Quantification of number of endolysosomal structures observed in ultrastructural analysis by TEM. For this analysis, MVEs were considered as endolysosomes as they frequently appeared as hybrid structures. Data are mean ± SD. The Student’s t-test was conducted with GraphPad Prism.

Noticeably, at higher magnification (25000x), we detected ELs approaching the plasma membrane (PM) in control cells compared to cytoplasmic accumulation of heterogeneous endolysosomes in the *SNCA*-KO cells (Supplementary Fig. S1A). Stage i and ii melanosomes were excluded from our definition used for quantifying endolysosomes. However, we observed a build-up of mostly stage ii melanosomes in *SNCA*-KO cells (Supplementary Fig S1B), which is consistent with a recent study on healthy melanocytes (32). Note that stage iii and iv melanosomes are indistinguishable from endolysosomes at the magnification we used for quantitative purpose due to similar electron density and hence, they were counted as endolysosomes.

Given that CD63 and LAMP1 reside on the limiting membrane of the endolysosomal organelles and that endolysosomes accumulate in the *SNCA*-KO cells (33, 34), we predicted a buildup of each of these proteins in KO compared to control cells. Strikingly, Western blot analysis of cell lysates from control and *SNCA*-KO cells revealed significant increases in the protein levels of CD63 (*P* < 0.0001) and LAMP1 (*P* = 0.0331) compared to control cells (Fig. 2A, B and C). As a further demonstration that the levels of CD63 and LAMP1 are sensitive to the amount of the α-syn protein, the levels of both CD63 and LAMP decreased (*P* < 0.0001, *P* = 0.0472, respectively) to the level of control cells upon lentiviral re-expression of wild type *SNCA* in *SNCA*-KO cells (labeled as *SNCA*-KI cells) (Fig. 2A, B and C). We also found a negligible increase in the levels of early endosome marker EEA1 in *SNCA*-KO cells, which was not statistically significant (*P* = 0.9238), and the level was variable in *SNCA*-KI cells, i.e., one biological replicate showed a level similar to the control while the other was increased (Fig. 2D). RT-qPCR analysis revealed no significant (*P* = 0.2951) change in the mRNA-levels of CD63, which confirmed that the large increase in the level of the CD63 protein in the *SNCA*-KO cells was not due to increased levels of mRNA (Fig. 2E). Taken together, there is an accumulation of endolysosomes but not early endosomes in *SNCA*-KO melanoma cells relative to control cells that express α-syn.

**Figure 2.**
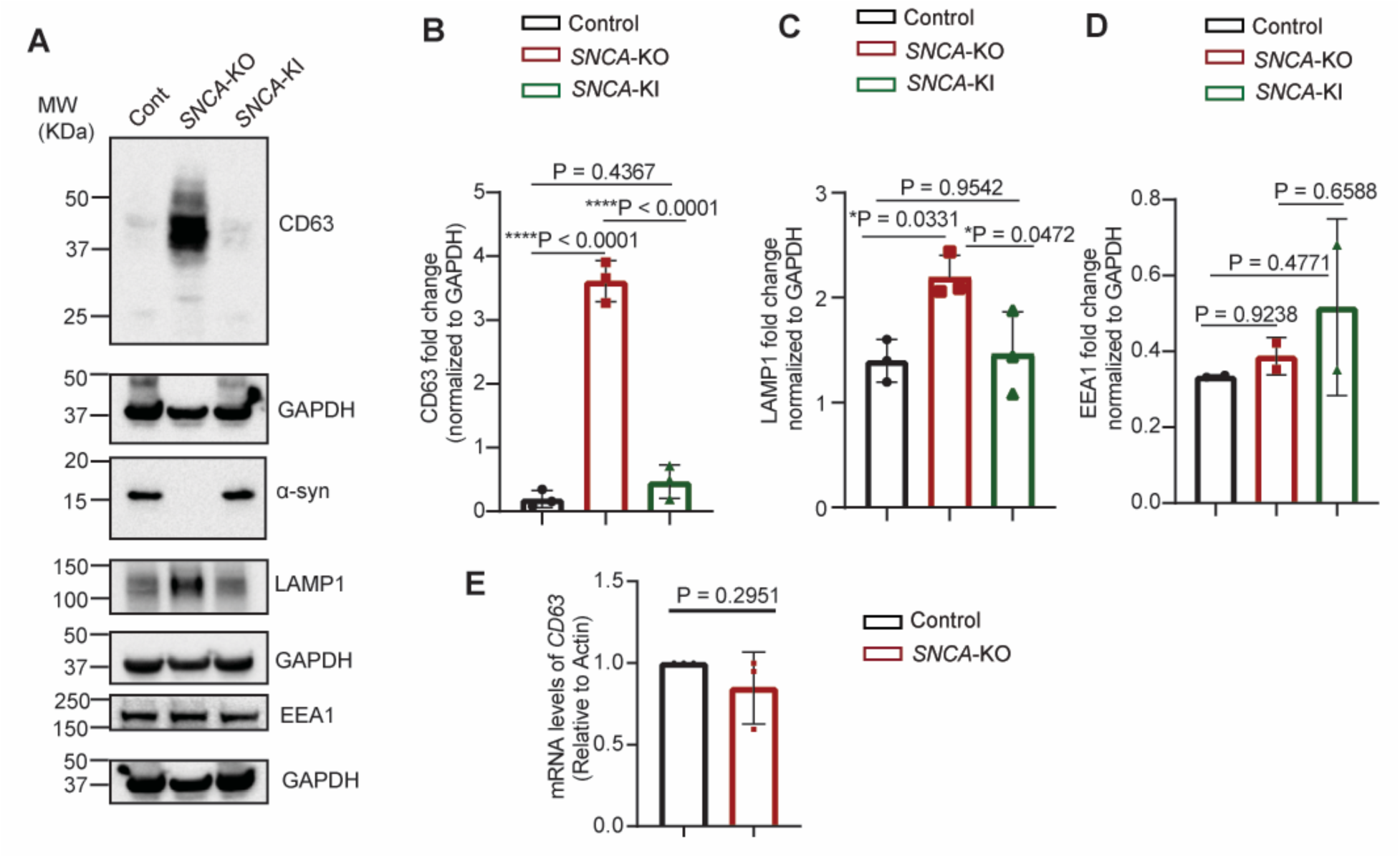
Endolysosomal membrane proteins are elevated in α-syn depleted cells. (A) Representative western blot images of endolysosomal membrane proteins (CD63, LAMP1). EEA1 was a marker for early endosomes. (B-D) Densitometric quantification of western blots shown in (A) for CD63 (B) LAMP1 (C), EEA1 (D). Fold changes were determined by normalizing band intensities to GAPDH. N = 3 per group for B-C and N=2 for D. Data are mean ± SD. For B-D, One-way ANOVA with Tukey’s correction was performed in GraphPad Prism. E. RT-qPCR showing the fold changes in mRNA level of *CD63* relative to actin. N = 3 per group. Data are mean ± SD. The Student’s t-test was conducted with GraphPad Prism. Uncropped blots are in Supplementary Fig. S2.

### Increased perinuclear clustering of endolysosomes occurs in α-syn-deficient melanoma cells

LAMP1 and CD63, which are found in both the limiting membrane (33, 34) and intraluminal vesicles (ILVs) of endolysosomes (33), are well-known endolysosome markers. We wanted to determine what portion of these two proteins were located on the limiting membrane of endolysosomes, and to address this we performed immunofluorescence to visualize these two proteins on endolysosomal membranes. We used 0.1% saponin, which selectively permeabilizes the cell membrane but preserves membranes of cytoplasmic organelles (35, 36). Immunolabeling for CD63 and LAMP1 showed localization to endolysosomal limiting membranes in control and *SNCA*-KO cells, and, at identical instrumental conditions, the fluorescence intensities of both CD63 and LAMP1 were markedly increased in *SNCA*-KO cells compared to control cells (Fig. 3). The distribution of both proteins appeared diffuse in the control cells compared to perinuclear clustering and colocalization in the *SNCA*-KO cells (yellowish region shown by arrowheads in Fig. 3). Often CD63 and LAMP1 colocalized to large juxtanuclear vacuoles in *SNCA*-KO cells (Fig. 3, inset in right middle panel and the lower right panel).

**Figure 3.**
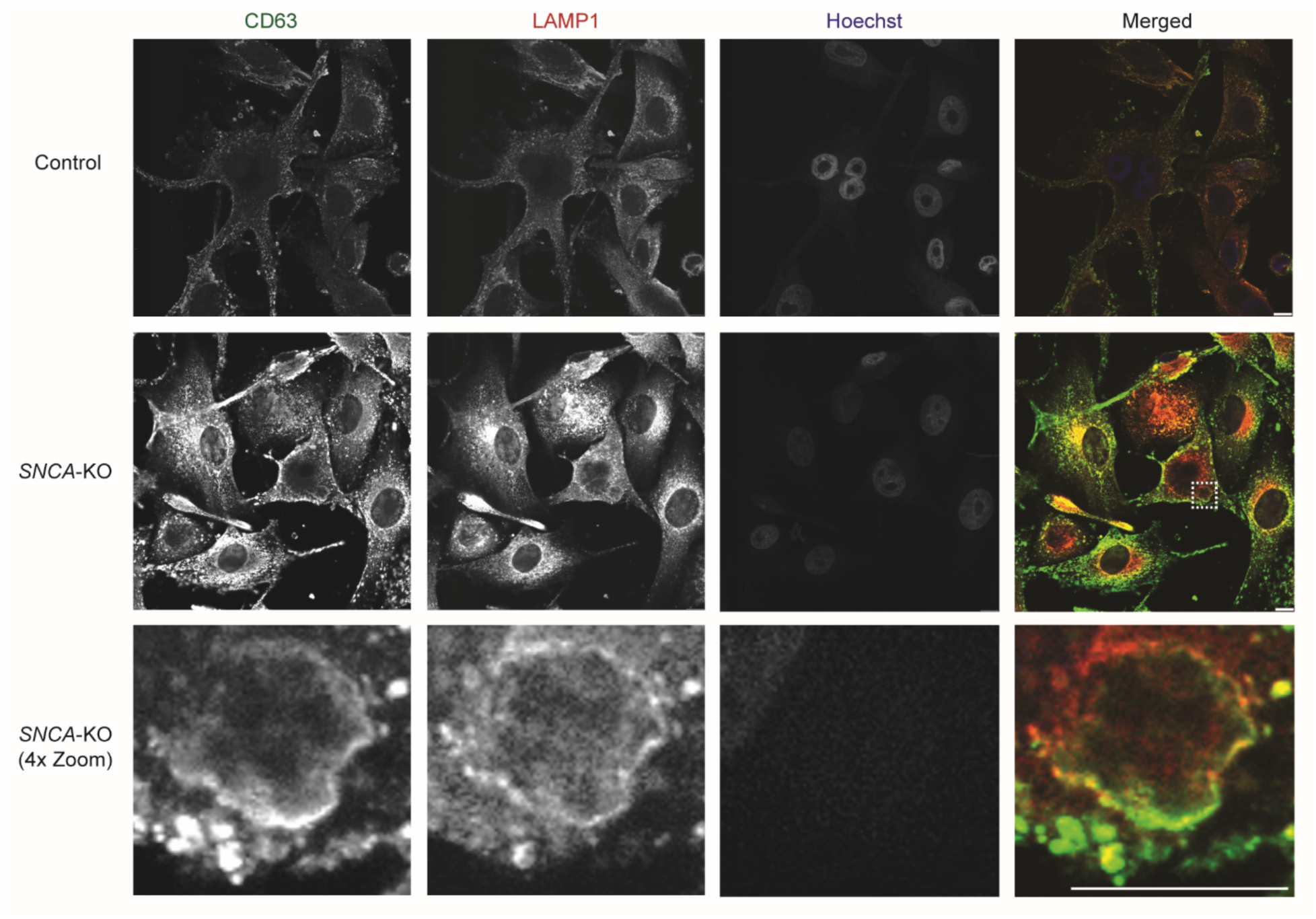
Endolysosomal membrane proteins cluster in large perinuclear vacuolar structure in SNCA-KO melanoma cells. Representative confocal microscopy images of immunostaining for CD63 (Green) and LAMP1 (Red) in control and *SNCA*-KO melanoma cells. Boxed area in the middle panel is enlarged (4x zoom in ImageJ) in insets in the lowest panel. Images are from the similar plane from Z-stacks. White arrowheads indicate colocalization of CD63 and LAMP1. Representative images from two independent experiments are shown. Scale bars = 10 μm.

We also measured the percentage of cells having large endolysosomes in control and *SNCA*-KO cells by TEM, where we defined a large endolysosome as having a diameter ≥ 800 nm. We found that *SNCA*-KO cells had a significantly higher percentage of cells (64.3%; *P* = 0.0124) with large endolysosomes (>800 nm) compared to control cells (40.8%) (Fig. 4A and B). We measured the distance of these large endolysosomes from the nuclear periphery, and it was revealed that the average distance of large endolysosomes from the nuclear periphery was significantly (*P* < 0.0001) shorter in *SNCA*-KO cells compared to the control cells. We did not observe a significant difference in the diameter of large (> 800 nm) endolysosomal organelles but interestingly endolysosomes with a diameter larger than 4 nm were only observed in *SNCA*-KO cells (Fig 4D). The results show that the perinuclear clustering of large endolysosomes is enhanced in α-syn deficient cells compared to control cells.

**Figure 4.**
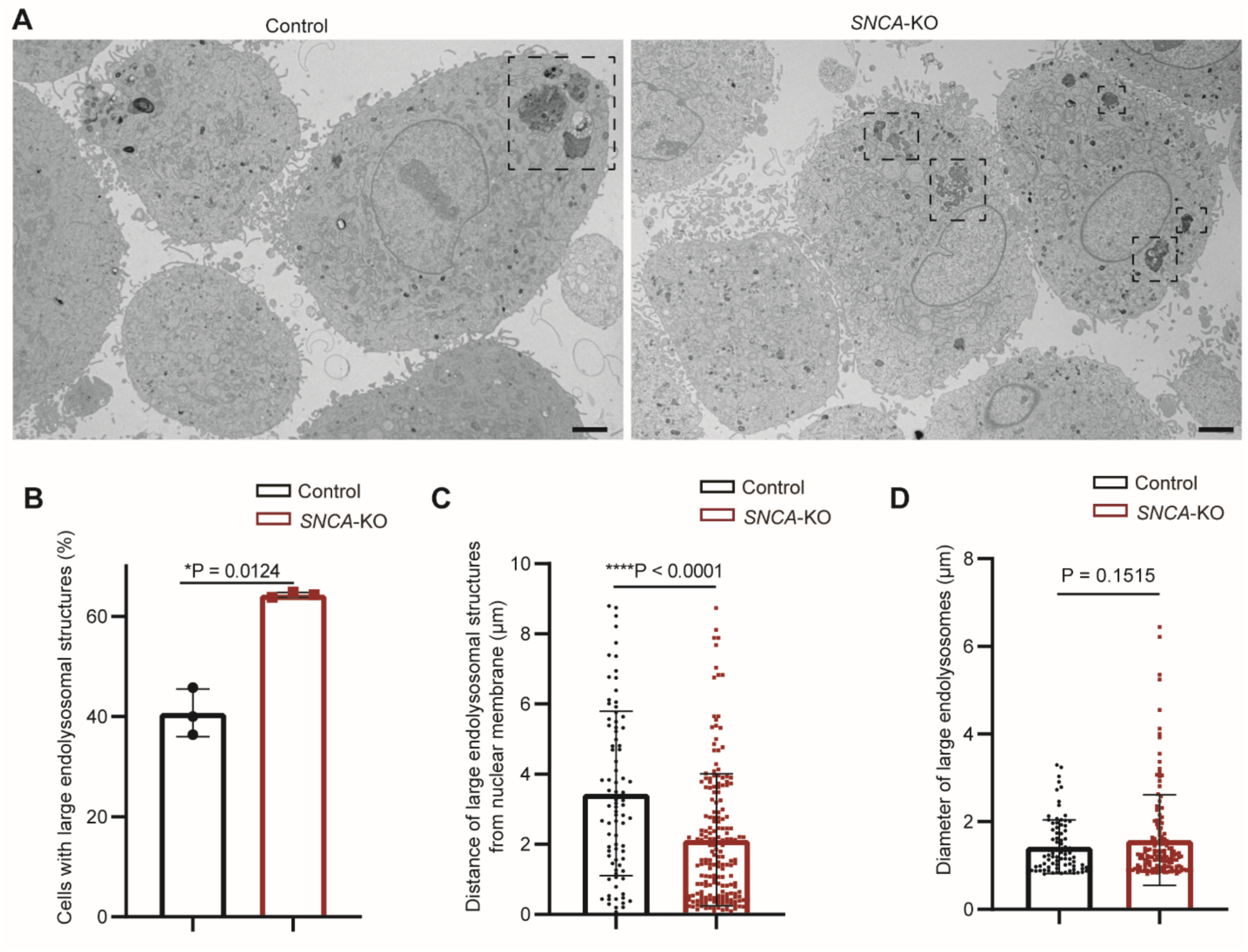
Large endolysosomal structures are more frequently observed in *SNCA*-KO cells with a perinuclear clustering pattern. A. Transmission electron microscopy images showing the pattern of clustering of large endolysosomal structures (defined as diameter > 800 nm) in Control vs *SNCA*-KO SK-MEL-28 cells. B. Percentage of cells with large endolysosomes (> 800 nm) in Control vs *SNCA*-KO cells. N = 3 for each group. C. Distance of edge of large endolysomal structures (> 800 nm) from the nuclear membrane (measured by ImageJ). D. Diameter of large endolysosomes in Control vs *SNCA*-KO cells. Data points in C-D indicate large endolysosomes. N = 141 cells (Control), N = 126 cells (*SNCA*-KO) were analyzed from three biological replicates. The Student’s t-test was conducted with GraphPad Prism.

### Pattern of crosstalk between heterogeneous endolysosomes in melanoma cells

We have shown that large endolysosomal organelles were more frequently observed in *SNCA*-KO cells compared to control cells (Fig. 4B). To gain insight into the fusion of heterogeneous endolysosomes, we observed the cells in higher magnification by TEM. An endolysosomal cluster containing multiple MVEs with clearly visible ILVs and endolysosomes with multilamellar and vesicular features are shown in *SNCA*-KO cells (Fig. 5, right panel, inset). In comparison, control cells mostly presented separate MVEs (Fig. 5, left panel). Distinguishing MVEs from amphisomes was not the objective of our study, however, a few structures with both ILVs and double membrane could be identified as amphisomes (Fig. 5, left inset). It should be noted that in earlier literature, the endolysosomal degradative compartments were often referred to as degradative autophagic vacuoles (aVd) (37, 38). The crosstalk between the autophagic compartments with endolysosomal compartments is well-known (39) but is not a part of our current study. Future studies are required to understand the complex crosstalk between autophagosomes (APs) and endolysosomes.

**Figure 5.**
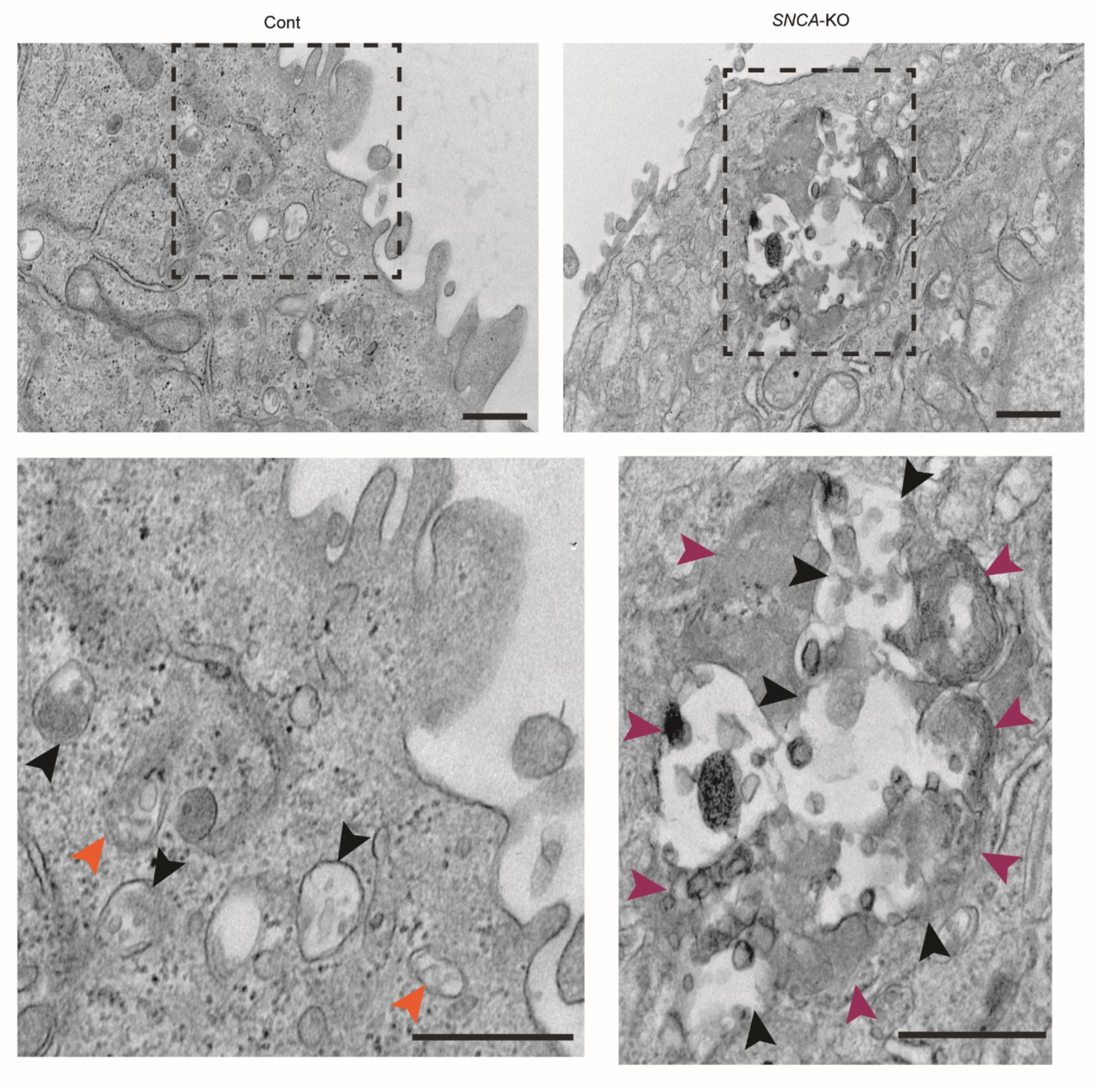
Endolysosomal fusion in melanoma cells. Ultrastructural analysis by transmission electron microscopy of control and *SNCA*-KO cells grown in polycarbonate membrane showing the clustering and fusion of multiple multivesicular endosomes (MVEs), amphisomes and endolysosomes. Boxed areas (dashed rectangles) from the corresponding cells are enlarged (1.5x zoomed in ImageJ) in the inset in the lower panel. Magenta arrowheads indicate endolysosomes (also includes multilamellar dense-core lysosomes shown in upper left and right parts of the inset for *SNCA*-KO group). Black arrowheads indicate MVEs (with less electron-dense intra-luminal vesicles compared to endolysosomes) and orange arrowheads indicate amphisomes (hybrid structures formed by fusion of autophagosomes and MVEs). Direct magnification = 10,000x. All Scale bars = 400 nm.

### CD81 and CD9 protein levels are reduced in *SNCA*-KO cells because of decreased expression and increased degradation

We recently reported a significant reduction in the level of the CD81 protein in *SNCA*-KO melanoma cells relative to control cells and found that some of the lower expression is due to a 40% reduction of *CD81* mRNA in the KO cells (27). We have previously shown enhanced lysosomal degradation of TFR1 (10) and L1CAM (26) in *SNCA*-KO cells, and here we asked whether the CD81 was also subject to enhanced lysosomal degradation in the KO cells. To test for endolysosomal degradation, control and *SNCA*-KO melanoma cells were treated with DMSO or Bafilomycin A1 (BafA, endolysosomal V-ATPase inhibitor) for 5 hours, and then cells were lysed and subjected to SDS PAGE and Western blotting. The level of CD81 was approximately 70% lower in the DMSO-treated *SNCA*-KO cells compared to control cells (Fig. 6B; compare columns 1 and 3). Treatment of *SNCA*-KO cells with BafA resulted in a significant (*P* = 0.0488) increase in the level of CD81 (Fig. 6B, compare columns 3 and 4), although the level of CD81 did not recover to that of the control cells. In contrast, treatment of the control cells with BafA had no significant effect on the level of CD81 (columns 1 and 2).

**Figure 6.**
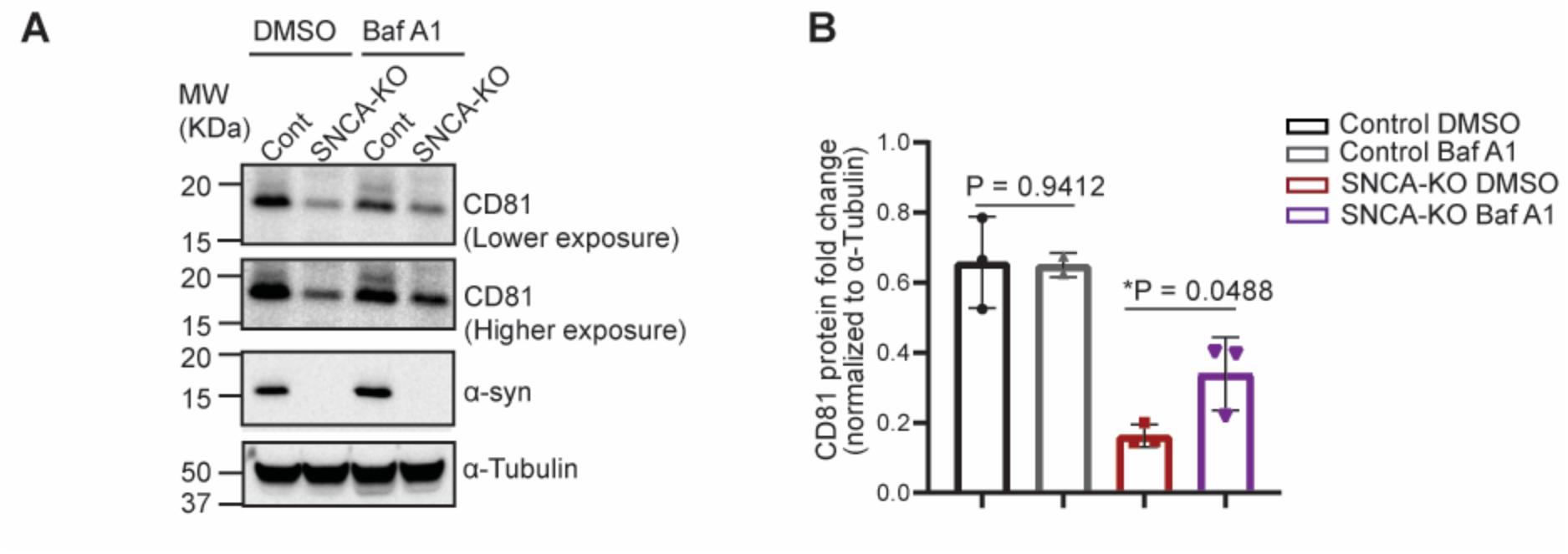
CD81 protein level is partially recovered upon inhibition of endolysosomal acidification suggesting a role of α-syn in CD81-trafficking. A. Western blots showing the level of CD81 in control and *SNCA*-KO cells treated either with DMSO (vehicle control) or Bafilomycin A1 (100 nM for 5 hours). B. Quantification of western blot images showing the fold change of CD81 protein normalized to α-Tubulin. N = 3 per group. Students t-test between the DMSO treated and BafA treated cells from each group were performed in Graphpad Prism. Data are mean ± SD. Uncropped blots are in Supplementary Fig. S3.

We also addressed whether knocking out of α-syn influenced the level of tetraspanin CD9. To this end, we observed a significant (*P* = 0.0007) decrease in the protein level of CD9 (band corresponds to CD9 isoform 2, NP_001317241.1 around ∼18 KDa) in *SNCA*-KO cells compared to the control cells, and the protein level was fully rescued (*P* = 0.0038) in *SNCA*-KI cells (lentiviral re-expression of SNCA into *SNCA*-KO cells) (Fig. 7A, B). We noticed multiple bands in these blots, which showed the presence of CD9-isoform 1 (∼26 KDa, Supplementary Fig. S4A). However, CD9-isoform1 also showed the same decreasing pattern, although that was not statistically significant (*P* = 0.0.852) (Supplementary Fig. S4B). It should be noted that the relative abundance of CD9-isoform 1 was much less than isoform-2 (Supplementary Fig. S4B). We then addressed whether CD9 was also undergoing endolysosomal degradation in *SNCA*-KO cells. We observed an elevation of CD9 (isoform-2) in *SNCA*-KO cells upon BafA treatment compared to DMSO-treated cells (Fig. 7C, D). However, this increase was not statistically significant (*P* = 0.3631) due to high error rates, but importantly, both independent experiments yielded an increase in CD9 upon BafA treatment (Fig. 7D). We also performed RT-qPCR, which revealed a significant reduction (*P* = 0.0361) in mRNA levels of CD9 (Fig. 7E). It should be noted that the CD9-isoform-1 was not detectable by the second antibody we used (Supplementary Fig. S4C, D). Collectively, the combined results indicate that the genetic depletion of α-syn causes a decrease in the level of both CD81 and CD9 protein, and these decreases are a consequence of a combination of enhanced endolysosomal degradation and decreased levels of their respective mRNAs.

**Figure 7.**
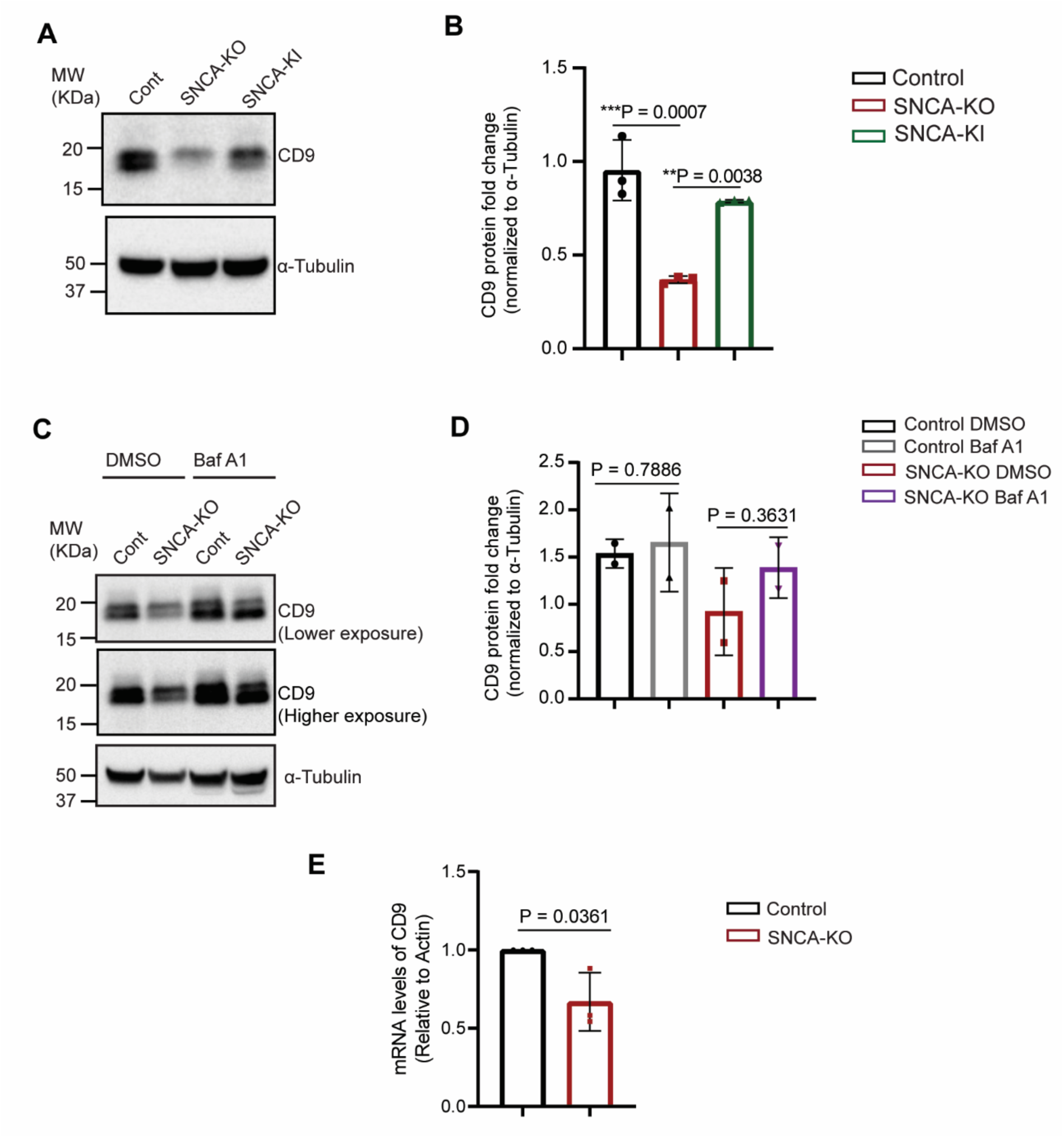
The expression and trafficking of tetraspanin CD9 is modulated by α-syn. A. Representative immunoblots showing a decrease in CD9 protein level in *SNCA*-KO cells, which were recovered upon re-expression of wild type α-syn (*SNCA*-KI cells). B. Quantification of immunoblots in A showing fold change of CD9 normalized to α-Tubulin. N = 3 per group. One-way ANOVA with Tukey’s correction was performed to calculate the P-values. C. Western blots showing the effect of BafA-treatment in protein level of CD9 in control and *SNCA*-KO cells. Cells were treated either with DMSO (vehicle control) or Bafilomycin A1 (100 nM for 5 hours). D. Quantification of western blot images showing the fold change of CD9 protein normalized to α-Tubulin. N = 2 per group. Student’s t-test between the DMSO treated and BafA treated cells were performed. E. RT-qPCR showing the mRNA fold change of *CD9* relative to actin. N = 3 per group. Data are mean ± SD. The Student’s t-test was conducted with Graphpad Prism. Uncropped blots are in Supplementary Fig. S4.

Previous studies suggested that both CD63 and CD9 were associated with promotion (40, 41) or inhibition (42) of melanoma progression in cell culture models. We asked whether expression of CD63 and CD9 were associated with patient survival. For this, we analyzed the TCGA-SKCM transcriptome dataset which revealed that patient survival was unaffected by the mRNA levels of CD63 or CD9 (Supplementary Fig. S5). Of the three tetraspanins (CD9, CD63, and CD81) we have studied in the context of melanoma, only CD81 affects patient survival (27).

## Discussion

The long-term goal of our work is to reveal the possible mechanism(s) for the involvement of α-syn in melanoma progression. This work was undertaken to gain insight into the mechanism by which loss of α-syn expression reduces the levels of many membrane proteins that are critical for invasion and signaling. The simplest possible explanation for the reduced levels of membrane proteins in our α-syn-deficient melanoma cells is that anterograde trafficking is compromised, which leads to vesicle accumulation and changes in lysosomal organelle positioning, plus there is less expression of proteins (due to reductions in their transcript levels and/or from degradation in the endolysosomes). These aspects are discussed below.

Loss of α-syn expression in SK-MEL-28 melanoma cells cause pathobiological changes in vesicle trafficking---the accumulation of endolysosomes and the positioning of large (> 800 nm) endolysosomes close to the nucleus relative to control cells—that, together with lysosomal protein degradation, are major contributing factors to the reduced levels of various membrane proteins in the SK-MEL-28 *SNCA*-KO cells. Overall, we propose that α-syn is an accessory factor that promotes the anterograde transport of endolysosomes and their cargo membrane proteins, as described in our model (Fig. 8). The essence of our model is that α-syn is a molecular tether: The 100 N-terminal residues can adopt an α-helical structure that avidly binds to membranes (43, 44), whereas the unstructured C-terminus (residues 101 – 140) can bind to proteins embedded in membranes. For example, α-syn tethers a MVE/Endolysosomes to the plasma membrane, as depicted in Fig. 8A (left panel), whereas loss of α-syn expression untethers the MVEs/Endolysosomes from the plasma membrane resulting, we propose, in a perinuclear clustering (Fig. 8A, right panel). α-syn can also tether an endolysosome to the kinesin machinery (Fig. 8B, left panel), and such a tethering function likely facilitates anterograde transport of endolysosomes. We propose that loss of α-syn expression slows down anterograde transport, resulting in the rate of retrograde transport dominating. An imbalance in anterograde/retrograde traffic upon the loss of α-syn expression explains the accumulation of large endolysosomal structures in the perinuclear region (Fig. 8A, right panel).

**Figure 8.**
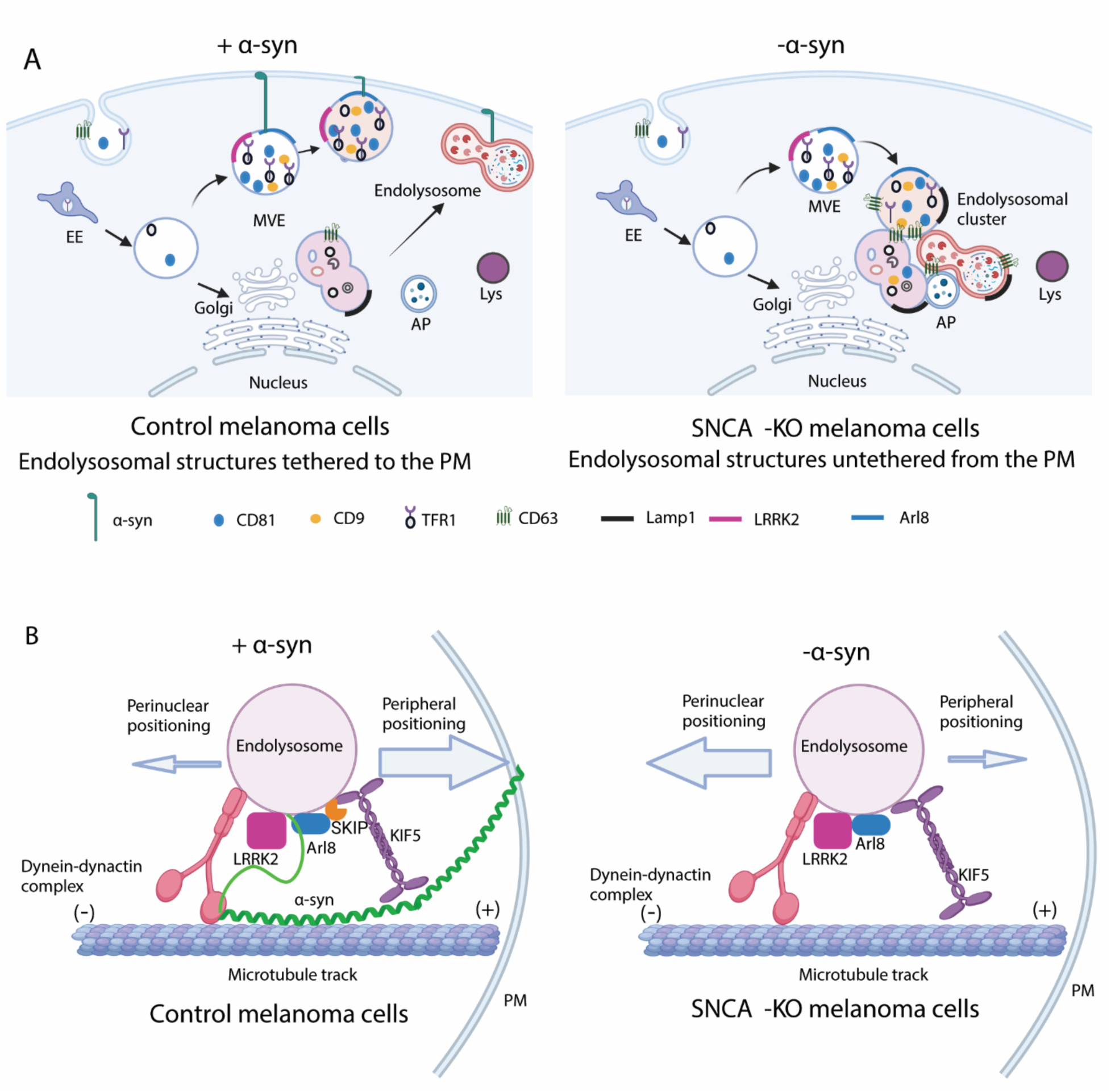
A model depicting the role of α-syn in modulation of positioning of endolysosomal structures. (A) α-syn tethers the endolysosomal structures to the plasma membrane (PM) in control melanoma cells (left sided panel). In absence of α-syn, endolysosomal structures are untethered from the PM and cluster in the perinuclear region of the cytoplasm. (B) A proposed detailed model of α-syn mediated modulation of trafficking of endolysosomal structures. We propose that in control melanoma cells, α-syn colocalizes with LRRK2 and ARL8 (54) on the MVE-membrane as reported in neuronal cells. The N-terminus of α-syn binds to the PM and α-syn is known to be associated with microtubules in melanoma cells. We propose that these dynamic interactions of α-syn with KIF5 and microtubules facilitate the anterograde trafficking of the endolysosomal structures to the plasma membrane (tethering) in control melanoma cells. In α-syn deficient melanoma cells, the retrograde trafficking of endolysosomal structures become dominant.

A wealth of data supports our model. For example, (i) α-syn binds to three families of proteins that are the main players in the transport and positioning of endolysosomes in mammalian cells (45), i.e., kinesins (46), the dynein-dynactin complex (47) and microtubules (48) in HEK293 cells and neuronal cells. (ii) α-syn binds to tubulin in both SK-MEL-28 and SK-MEL-5 melanoma cells (49). (iii) α-syn and syb2/VAMP2 colocalize in neuronal vesicles (15), and α-syn colocalizes with VAMP2 at exocytic cites on unroofed SK-MEL-28 cells (50). This finding suggests that α-syn/VAMP2 colocalization can happen in other vesicular compartments than neuronal vesicles in other mammalian cells. (iv) α-syn interacts with LC3-II on the MVB/endomembrane compartments in induced pluripotent stem cell (iPSC) derived neuronal cells (51). (v) α-syn interacts with CHMP2B, another MVE-protein belonging to the Endosomal Sorting Complex required for Transport III (ESCRT-III) family (52). (vi) Arl8, a key protein involved in movement and positioning of endolysosomes (45, 53), colocalizes with LRRK2 and α-syn (54).

A straightforward way to test this model is to re-introduce a variant that lacks the tether (α-syn[1-100]) into the SK-MEL-28 *SNCA*-KO cells. Our prediction is that *SNCA*-KO cells expressing α-syn[1-100] would phenocopy the *SNCA*-KO cells. On the other hand, C-terminally truncated variants of α-syn are toxic to neurons (55) and thus might be also toxic to the melanoma cells, so SK-MEL-28 cells that express this variant might not behave exactly like the KO cells. This is a future direction.

In SK-MEL-28 *SNCA*-KO cells, we have found significantly reduced levels of TFR1, L1CAM, N-Cadherin, FPN1, CD9 and CD81, and that for three (TFR1, CD9, and CD81) of these proteins their respective mRNA levels were also significantly reduced (10, 26, 27) (Table 1). The mRNA level of L1CAM was slightly higher (but not significantly) in the *SNCA*-KO cells, and mRNA levels for N-Cadherin and FPN1 were not determined. Ben Gadalya and colleagues also reported a positive correlation between α-syn protein level and TFR1 protein level in MN9D neuronal cells (21). Our interpretation is that the reduction in TFR1, CD9, and CD81 protein levels in the *SNCA*-KO cells is partly from lower expression due to decreased levels of mRNA (relative to control cells). We do not know why mRNA levels decrease for these three proteins in the SK-MEL-28 *SNCA*-KO cells. Given that loss of α-syn expression in SK-MEL-28 cells results in the significant downregulation of 660 transcripts (28), the transcripts for TFR1, CD9, and CD81 might be reduced in a homeostatic response to changes in transcript levels of genes involved in numerous pathways. Alternatively, perhaps in control cells α-syn enters the nucleus and plays a role in transcription of these genes or DNA repair (56, 57) or it promotes the processing, transport or stability of mRNAs (58); consequently, upon knocking out *SNCA* these transcriptional or RNA processing functions are lost, resulting in decreases in transcript levels. Overall, the reduced levels of some membrane proteins in the SK-MEL-28 *SNCA*-KO cells are partly due to lower expression.

**Table 1.** Membrane proteins affected by the loss of α-syn expression in SK-MEL-28 cells.

| protein | Protein level | mRNA | Partial Lysosomal degradation | references |
| --- | --- | --- | --- | --- |
| TFR1 | ↓ | ↓ | yes | (21) |
| CD9 | ↓ | ↓ | yes | This study |
| CD81 | ↓ | ↓ | yes | (27), this study |
| N-Cadherin | ↓ | ND | ND | (26) |
| L1CAM | ↓ | NC | yes | (26) |
| FPN1 | ↓ | ND |  | (21) |
| CD63 | ↑ | NC | ND | This study |
| Lamp1 | ↑ | ND | ND | This study |

NC, no change; ND, not determined.

In melanoma cells with elevated expression of α-syn, there is robust anterograde membrane traffic that results in the delivery of key adhesion molecules, which control invasion and migration, to the plasma membrane, i.e., L1CAM and CD81 and others. Knocking out *SNCA* is devastating because in the absence of α-syn L1CAM and CD81 and other key proteins involved in metastasis mistraffic and are degraded in the lysosome. The result is that such low α-syn-expressing cells should be less aggressive than melanoma cells with elevated expression of α-syn. This idea is borne out by our previously published work showing that individuals with melanomas with expression of both α-syn of CD81 have significantly diminished survival (27). As we have previously demonstrated that SK-MEL-28 melanoma cells expressing SNCA were more invasive (26), our current model supports the idea that peripheral positioning promotes invasiveness by promoting endolysosomal secretion of proteases and EVs in cancer cells as reported in previous studies (59, 60), and reviewed in (61).

In the future, targeting *SNCA* mRNA with an antisense oligonucleotide (ASO) delivered by tissue specific adenoviral vector together with other chemotherapeutic agents that are currently used for melanoma could be a novel strategy to suppress melanoma metastases by lowering the levels of the key adhesion and signaling molecules, L1CAM and CD81. Using an ASO might be better than targeting L1CAM and CD81 with antibodies, as two monoclonal antibodies would have to be used, which would likely be toxic to melanoma patients. Future work is required to understand the mechanistic and structural details of how α-syn interacts with the molecular machinery involved in endolysosomal transportation.

## Materials and Methods

### Cell culture

SK-MEL-28 cells were purchased from ATCC. The gene *SNCA* that codes for α-syn was knocked out in SK-MEL-28 cells using CRISPR/Cas9 and characterized in detail (10); these knockouts (KO) are referred to as SK-MEL-28 *SNCA*-KO cells. Cells were cultured in DMEM supplemented with 10% FBS and 1% penicillin/ streptomycin at 37°C in 5% CO2. Cells were subcultured at 70-80% confluency.

### Transmission Electron Microscopy

For imaging adherent cells in situ, cells were grown either on Thermanox coverslips (Electron Microscopy Sciences, #174934) or on Nunc Cell Culture Inserts (Electron Microscopy Sciences, #72296-22) in six well plates in complete DMEM for 48 hours. Then, the cells were washed thrice with phosphate buffered saline (PBS, pH 7.4)) (Thermofisher, Gibco ^TM^, #10010023) and fixed in 2.5% glutaraldehyde + 2.5% formaldehyde in 0.1 M sodium cacodylate buffer (pH 7.4) (Electron Microscopy Science # 15949) for 2 hours in room temperature. The coverslips/polycarbonate filters were inspected under light microscope and 1 mm x 1 mm small pieces were cut. The post-fixation was done in 1% osmium tetroxide (OsO4) for 45 minutes followed by serial dehydration in graded acetone (50% for 6 minutes, 70% for 6 minutes, 95% for 10 minutes and 100% for 15 minutes). The samples were then infiltrated with a mixture of 100% acetone and increasing amount of 100% resin (2:1, 1:1, 1:2 and finally only in 100% Spurr resin). Finally, the samples were embedded in Spurr resin and polymerized at 65°C overnight. The images were obtained in a JEOL −1400 Flash transmission electron microscope.

For imaging sections of cell pellets, cells were grown for 48 hours in the abovementioned method and then washed thrice with PBS. Then, cells were gently detached from the plate with ethylenediaminetetraacetic acid (EDTA) (10 minutes at 37°C). Cells were pelleted by centrifuging at 3000 rpm for 3 minutes. The supernatant was carefully discarded, and the cell pellets were fixed in the abovementioned fixative for 2 minutes. Then, the pellets with the fixative were centrifuged again with great caution so that the cells were not resuspended in the fixative to allow proper fixation and crosslinking of cells. Cell pellets were fixed at room temperature for 2 hours. The fixed pellets were shipped to Harvard Electron Microscopy facility. Cells were washed in 0.1M cacodylate buffer and postfixed with 1% Osmium tetroxide (OsO4)/1.5% (K_4_FeCN_6_) for 1 hour, washed 2x in water, 1x maleate buffer (MB) 1x and incubated in 1% uranyl acetate in MB for 1hr followed by 2 washes in water and subsequent dehydration in grades of alcohol (10min each; 50%, 70%, 90%, 2×10 min 100%). The samples were then put in propyleneoxide for 1 hour and infiltrated overnight in a 1:1 mixture of propyleneoxide and TAAB Epon (TAAB Laboratories Equipment Ltd). The following day, the samples were embedded in TAAB Epon and polymerized at 60 degrees C for 48 hrs. Ultrathin sections (80 nm) were cut on a Reichert Ultracut-S microtome, picked up on to copper grids, stained with lead citrate. Images were acquired in a JEOL 1200EX transmission electron microscope. Images were recorded with an AMT 2k CCD camera.

### Confocal microscopy

Cells were grown in complete DMEM for 24 hours on cover glasses in six well plates. Then, the cells were washed thrice in PBS and fixed with 4% paraformaldehyde (Thermofisher # J61899.AP) for 15 minutes. Then, cells were permeabilized with 0.1% saponin (Sigma-Aldrich #SAE0073-10G) for 10 minutes followed by 60 minutes blocking in 5% goat serum. Cells were incubated in mouse and rabbit primary antibodies (diluted in 5% goat serum in 0.1% saponin) in a humidified box overnight at 4°C. Then, after 3 washes (2 minutes for each wash) in ice cold PBS, incubation in secondary antibodies was performed at room temperature for 60 minutes. After washing the cells thrice in PBS, cells were incubated in Hoechst 33342 (Thermofisher, #H3570) to stain the nucleus. The cells were washed twice in PBS and the coverslips were mounted in ProLong^TM^ Glass Antifade Mountant (Thermofisher, #P36982). Images were acquired in the same laser power, gain, offset and aperture setting in Z-stacks (step size = 0.4 μm, voxel depth = 395.2) with the 100X objective of Leica TCS5 confocal microscope. Sequential scanning by frame (In the order of channels for 488 nm, 594nm and 405 nm) were performed at 400 Hz.

### Image analysis

Images were acquired using the Leica LAS software (version 1.4.6). Confocal images were processed in ImageJ software to obtain the zoom and grayscale image for each channel. There was no further editing done and each image is uncropped and showing the entire field. All the TEM images were unedited and uncropped. The entire field was shown for each TEM image. Measurements were done in ImageJ. Images were assembled in Adobe illustrator for final figures.

### Western blotting

Cells were grown in the above-mentioned conditions for 48 hours. After two washes with ice-cold PBS, cells were lysed in ice-cold RIPA buffer, and after passing through 25G needles with insulin syringes, the cell lysates were centrifuged at 18,360 g at 4°C for 20 minutes. Then, protein concentration was determined with BCA. Equal amount of protein was loaded into each lane of NuPAGE^TM^4-12% Bis-Tris Gel (Invitrogen by Thermofisher, #NP0335BOX, #NP0336BOX). The electrophoresis was performed at constant voltage, and the protein was transferred to nitrocellulose membranes (Biorad transblot Turbo Mini NC transfer packs, #1704158) using semi-dry transfer in Bio-Rad Trans-Blot Turbo Transfer System. Membranes were incubated in a 5% membrane-blocking agent (G-Bioscience, #786-011) for 1 hour at room temperature and then incubated with the primary antibody overnight at 4°C. Membranes were washed five times with 1X Phosphate Buffered Saline with 0.1%(v/v) Tween-20 (PBST) and incubated with HRP-conjugated secondary antibody for 1 hour at room temperature. Finally, bands were developed with Clarity ^TM^ western ECL substrate (Bio-Rad, #1705060) in a Bio-Rad ChemiDoc Touch machine. Images were obtained at different exposures. Membranes were cut, and the strips were re-probed for loading control.

### RT-qPCR

SK-MEL-28 cells were cultured for 48 hours at 37°C, 5% CO_2_ in complete DMEM medium. Total RNA was extracted using E.Z.N.A total RNA kit I (Omega Boi-Tek, #R6834-02) following manufacturer’s protocol. RNA quality and concentration was assessed with NanoDrop 2000c (Thermo Scientific, # ND2000CLAPTOP). cDNA was synthesized from 1μg of total RNA from each sample as follows: 1μg total RNA + 2 μL 5X first strand buffer + 1 μL DNAseI + DEPC treated ddH2O to make 10μL total volume. This was incubated for 30 minutes at 37°C. Then, 1 μL of DNAseI inhibitor was added and incubated at 65°C for 10 minutes and quenched on ice for 1 minute. The reverse transcription reaction was set up as follows: 5 μL DNAse treated RNA+1 μL 5X RT buffer + 0.25 μL random primers (1:30 dilution) + 0.25 μL dNTPs + 0.5 μL DTT + 0.2 μL RNAse-inhibitor + 0.2 μL MMLV RTase (Promega, #M1708) or (ddH20 for RT-) + 2.6 μL DEPC treated ddH2O = 10 μL final volume of mastermix was incubated at 37°C for 1 hour. This reaction was set up for RT- and no primer control for each sample to control for any non-specific amplification in qPCR. Finally, the samples were treated with RNAseH at 37°C for 30 minutes. qPCR was performed using the iTaq Universal SYBR Green Supermix (Bio-Rad, Cat. #1725121) according to the manufacturer’s protocol in a Bio-Rad CFX Opus 96 Real-Time PCR System (Bio-Rad, #12011319). Relative fold changes were analyzed in 2^-ΔΔCT^ method. Primers used:

CD63: Forward: 5’-CAACGAGAAGGCGATCCATAA-3’

Reverse: 5’-GCAGCAGGCAAAGACAATTC-3’

CD9: Forward: 5’-CTGTTCGGCTTCCTCTT-3’

Reverse: 5’-CTCCTGGACTTCCTTAATCACC-3’

Actin: Forward: 5’-GGAGAAGAGCTACGAGCTGCCTGAC-3’

Reverse: 5’-AAGGTAGTTTCGTGGATGCCACAGG-3’

Primers were ordered from Integrated DNA Technologies.

### Antibodies

Mouse monoclonal antibodies: Human anti-CD63 (TS63, abcam, #ab 59479, 1:2000 for WB, 1:200 for IF); Anti-CD81 (B-11, Santa Cruz Biotechnolgy, #sc-166029, 1:500 for WB). Anti-CD9 (C-4, Santa Cruz Biotechnolgy, #sc-13118, 1:500 for WB). Anti-GAPDH (Novus Bio, # NB300-221, 1: 2000 for WB); α-Tubulin (Sigma Aldrich, #T9026, 1:2000 for WB). Rabbit monoclonal antibodies: CD9 (D3H4P, Cell Signaling Technology, #13403S). Rabbit polyclonal antibody: Anti-LAMP1 (Abcam, # ab24170, 1:1000 for WB, 1:100 for IF)

The horseradish peroxidase (HRP) conjugated secondary antibodies: Mouse (Santa Cruz Biotechnology, # sc–516102, 1:1000 or 1:2000 for WB) and Rabbit (1:2000, Santa Cruz Biotechnology, # sc–2357, 1:1000 or 1:2000 for WB). Fluorophore-conjugated secondary antibodies for immunofluorescence (IF): Goat anti-Mouse IgG (H+L) secondary antibody conjugated with Alexa Fluor^TM^ 488 (Invitrogen, #A11029, 1:200); Goat anti-Rabbit IgG (H+L) secondary antibody conjugated with Alexa Fluor^TM^ 594 (Invitrogen, #A11012, 1:200).

### Statistical analysis and software

Statistical analyses were performed using GraphPad Prism v8 (GraphPad Software, San Diego, CA; www.graphpad.com). A one-way analysis of variance (ANOVA) with a Dunnett post hoc test when comparing multiple groups. A two-tailed Student’s t-test was used when comparing two groups, assuming equal variances. A *P*-value ≤ 0.05 was considered significant. All data were expressed as means ± SD. The flow cytometry data were analyzed using the Sasquatch (SQS) software.

## Acknowledgements

This research was supported in part by several intramural grants from the Feist-Weiller Cancer Center (to SNW). We thank Dr. Brent Reed, Dr. Kevin McCarthy and Dr. Martin Muggeridge for their advice on confocal microscopy. We thank Dr. Andrew Yurochko for the gift of the EEA1 antibody and the actin primer. We thank Dr. Judy King for her advice on electron microscopy. We thank the electron microscopy facility of Department of Pathology and the research core facility for the microscope and the PCR instruments.

## Abbreviations

ASO: antisense oligonucleotide
Bafilomycin A1: BafA, endolysosomal V-ATPase inhibitor
DA: dopamine; dopaminergic
ILVs: intraluminal vesicles
MVEs: multivesicular endosomes
PD: Parkinson’s disease
SNARE: soluble *N*-ethylmaleimide-sensitive factor attachment protein receptor
SD: standard deviation
TCGA-SKCM: 
TEM: transmission electron microscopy
TFR1: transferrin receptor 1

