## Supplementary Figures 1-5. for "Alpha-synuclein modulates the positioning of endolysosomes in melanoma cells"

[1]Department of Biochemistry and Molecular Biology, Louisiana State University Health Sciences Center, 1501 Kings Highway, Shreveport, LA 71103, United States. [2]Feist-Weiller Cancer Center, LSU Health, 1405 Kings Highway, Shreveport, LA 71103, United States. [3]Harvard Medical School, Electron Microscopy Facility, 220 Longwood Ave., Boston, MA 02115, United States. [4]Ochsner-LSU Health, 1541 Kings Highway, Shreveport, LA 71103, United States.

This document contains Supplementary figures S1-5 and legends.

### Legends

**Figure S1.** Transmission electron microscopy images showing the accumulation of heterogeneous endolysosomal structures in *SNCA*-KO melanoma cells. (A) Upper panel: Accumulation of heterogeneous endolysosomal structures is the hallmark of *SNCA*-KO melanoma cells (right side) compared to the control cells (left side). Magnification = 8000x, scale bar = 500 nm. Lower panel left side: endolysosomal structures approximating the plasma membrane, which might lead into possible fusion events. Lower panel, right side: a higher magnification (not a zoom) view of the boxed region of the image in upper right panel, showing the clustering of heterogeneous endolysosomal structures. Magnification = 25000x, scale bar = 100 nm. (B) Melanosomes accumulate in *SNCA*-KO cells. TEM images showing accumulation of stage ii and stage iii melanosomes in *SNCA*-KO cells (right side) compared to the control cells (left side). Magnification = 25000x, scale bar = 100 nm. Arrowheads used in (A-B): Yellow (AP), orange (amphisome), blue (EE), black (MVE), and magenta (endolysosomes/autolysosomes). Representative images of three biological replicates are shown. These samples were prepared from ultrathin sections prepared from cell pellets as detailed in the method section.

**Figure S2**. (A-D). Uncropped and uncontrasted WB images shown in Figure 2A. (A) CD63 and the corresponding GAPDH reprobes. (B) LAMP1 and corresponding GAPDH reprobes. (C) α-syn, (D) EEA1 reprobed on the blot for the third biological replicate of the CD63 blot. So, these two blots had the same loading control (GAPDH). All the biological replicates were shown on the right side of the corresponding panel. Asterisks indicate non-specific bands. The solid rectangles represent the edges of the full-length blots. Blots containing the colorimetric image with the molecular ladders were shown wherever the edges were not clearly visible (right side of corresponding blots). Area boxed by dashed rectangles were shown in Figure 2A.

**Figure S3.** Uncropped and uncontrasted WB images shown in Figure 6A. (A-C) All biological replicates of CD81 immunoblotting in lower and higher exposures. Corresponding loading control (α-Tubulin) for each blot was shown in each panel. The solid rectangles represent the edges of the full-length blots. Area boxed by dashed rectangles were shown in Figure 6A.

**Figure S4.** Uncropped and uncontrasted WB images shown in Figure 7A, 7C. (A) Uncropped and uncontrasted immunoblots for Figure 9A from all biological replicates. Colorimetric merged image (at the right side) clearly shows the molecular ladders and blot edges. The solid rectangles represent the edges of the full-length blots. Area boxed by dashed rectangles were shown in Figure 7A. (B) Quantification CD9-isoform-1 (~26 KDa) detectable only with the Santa Cruz antibody. It should be noted that this isoform was not detectable with the other antibody (Cell Signaling Technology). (C-D) Uncropped and uncontrasted blots shown in Figure 7C from all biological replicates. The solid rectangles represent the edges of the full-length blots. Area boxed by dashed rectangles were shown in Figure 7C. Panel C (right side) shows the colorimetric image showing the molecular ladders. The loading control for C is the same as the loading control of Figure S3B (Biological replicate of CD81 expression on BafA treatment) because this same blot was reprobed for CD9.

**Figure S5.** Melanoma-specific survival probability is not affected in patients with high expression of *CD63* and *CD9* genes. (A) Melanoma-specific survival is unchanged in patients with higher expression of *CD63*. Red line depicts melanoma patients with higher expression of *CD63* (Exp > 0.5 SD above the median expression). Blue line depicts melanoma patients with lower expression of *CD63* (below above defined range). N = 420, Low expression group = 320, high expression group= 100. (B) Melanoma-specific survival is unchanged in patients with higher expression of *CD9*. The red line depicts melanoma patients with high expression (Exp > 0.5 SD above the median expression) of *CD9*. Blue line depicts melanoma patients with lower expression of CD9 (below the abovementioned range) N = 420, Low expression group = 312, high expression group= 108.  Z threshold = $\pm$ 0.5. The total number of patients in the analysis, N = 420. Disease-specific survival option was chosen from cBioportal to generate K-M survival curve.

Figure S1

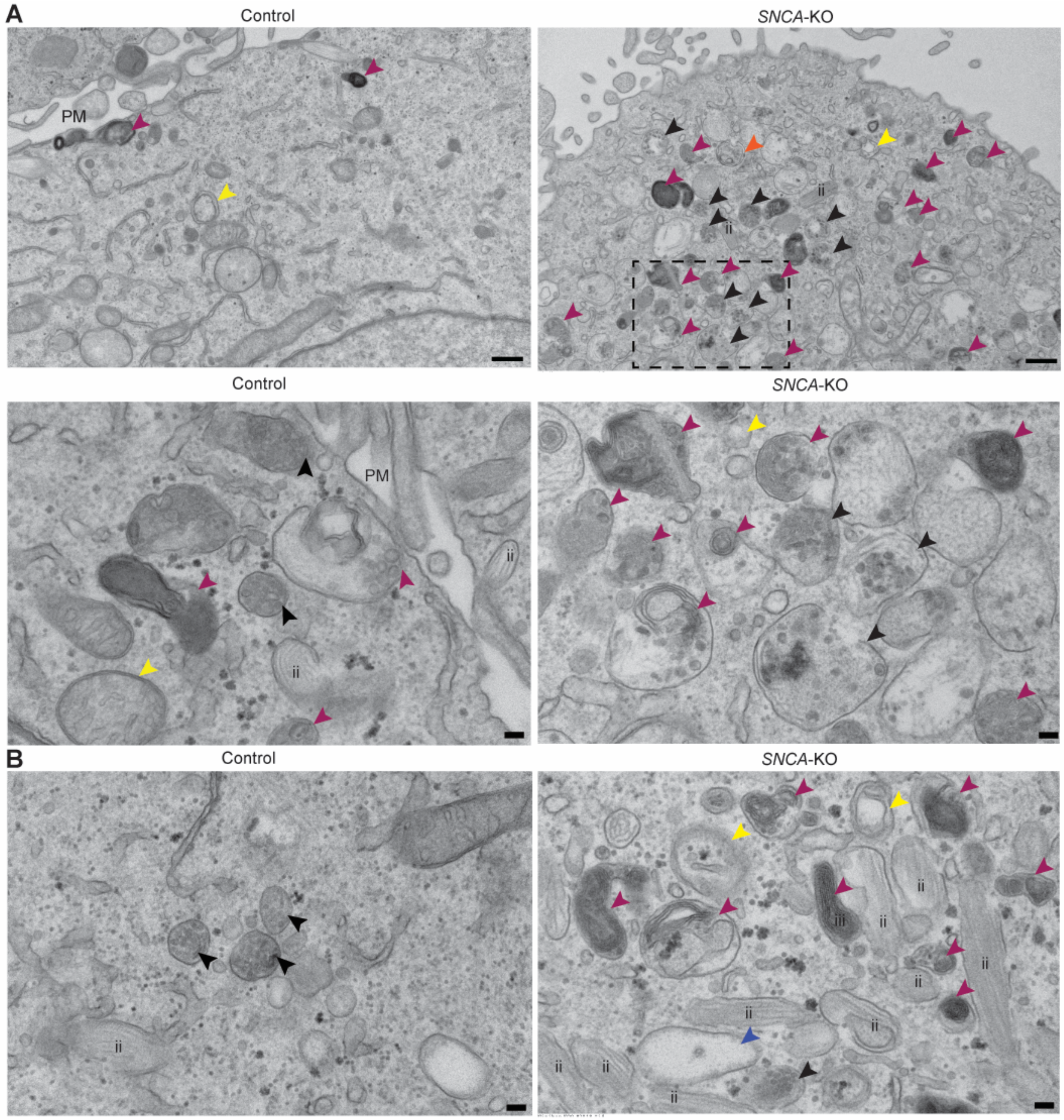
A
Control
SNCA-KO
PM
ii
Control
SNCA-KO
PM
ii
ii
B
Control
SNCA-KO
ii
ii
iii
ii
ii
ii
ii
ii
ii
ii
ii

Figure S2

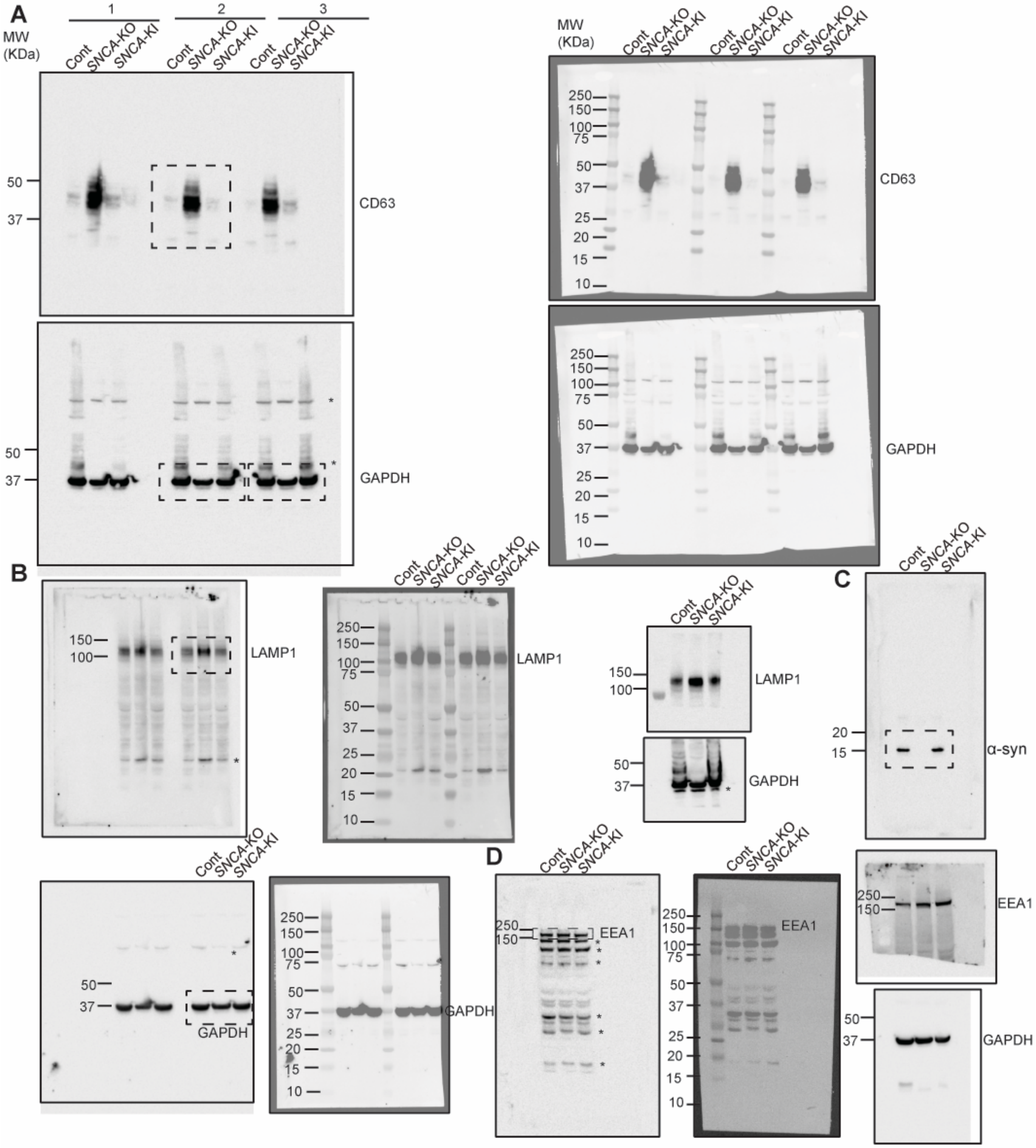

A
B
C
D
1
2
3
MW (KDa)
Cont
SNCA-KO
SNCA-KI
250
150
100
75
50
37
25
20
15
10
CD63
GAPDH
LAMP1
α-syn
EEA1
*

Figure S3

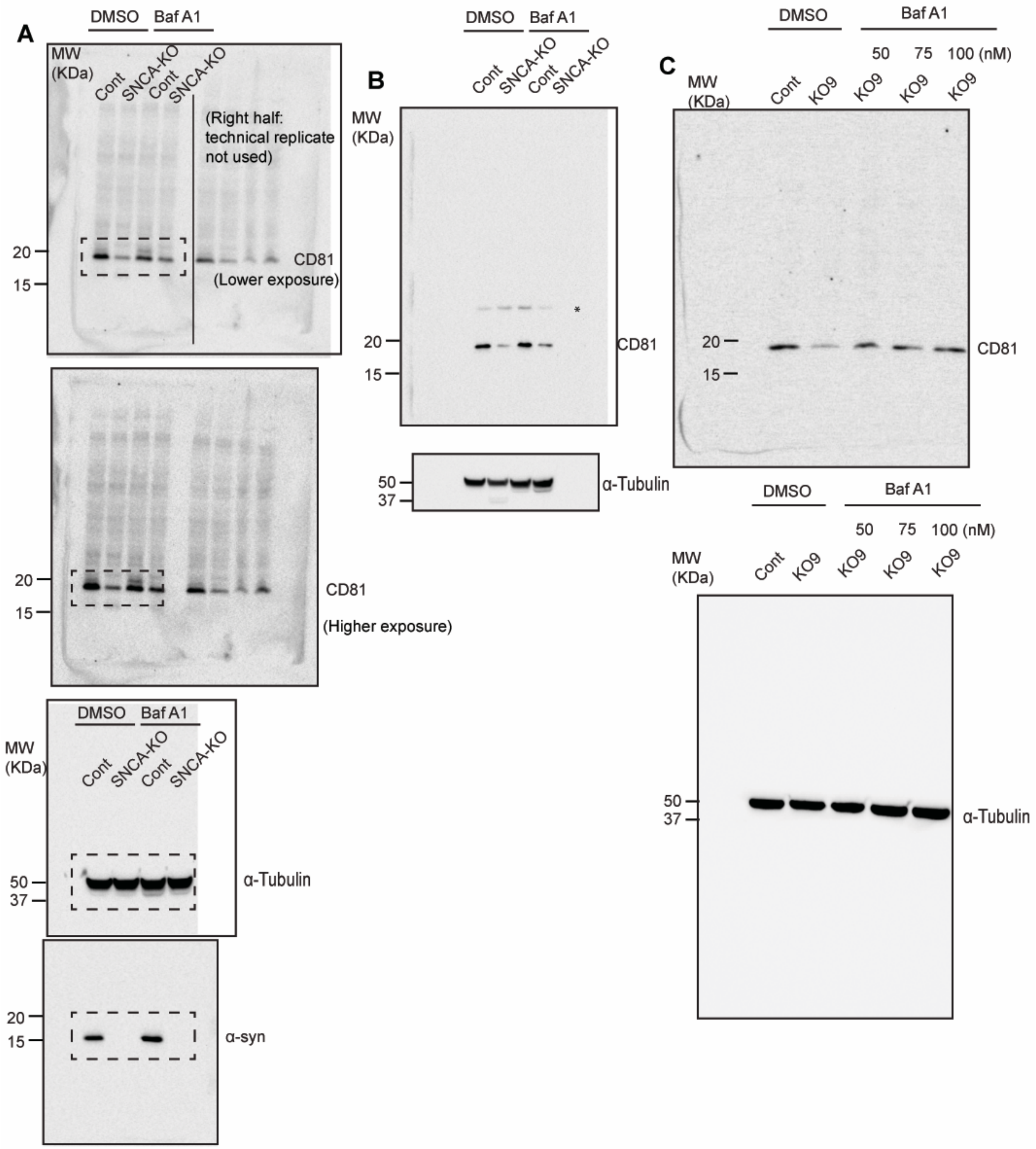
A
DMSO
Baf A1
MW (KDa)
Cont
SNCA-KO
Cont
SNCA-KO
(Right half: technical replicate not used)
20
15
CD81
(Lower exposure)
20
15
CD81
(Higher exposure)
DMSO
Baf A1
MW (KDa)
Cont
SNCA-KO
Cont
SNCA-KO
50
37
α-Tubulin
20
15
α-syn
B
DMSO
Baf A1
Cont
SNCA-KO
Cont
SNCA-KO
MW (KDa)
*
20
15
CD81
50
37
α-Tubulin
C
DMSO
Baf A1
50
75
100 (nM)
MW (KDa)
Cont
KO9
KO9
KO9
KO9
20
15
CD81
DMSO
Baf A1
50
75
100 (nM)
MW (KDa)
Cont
KO9
KO9
KO9
KO9
50
37
α-Tubulin

Figure S4

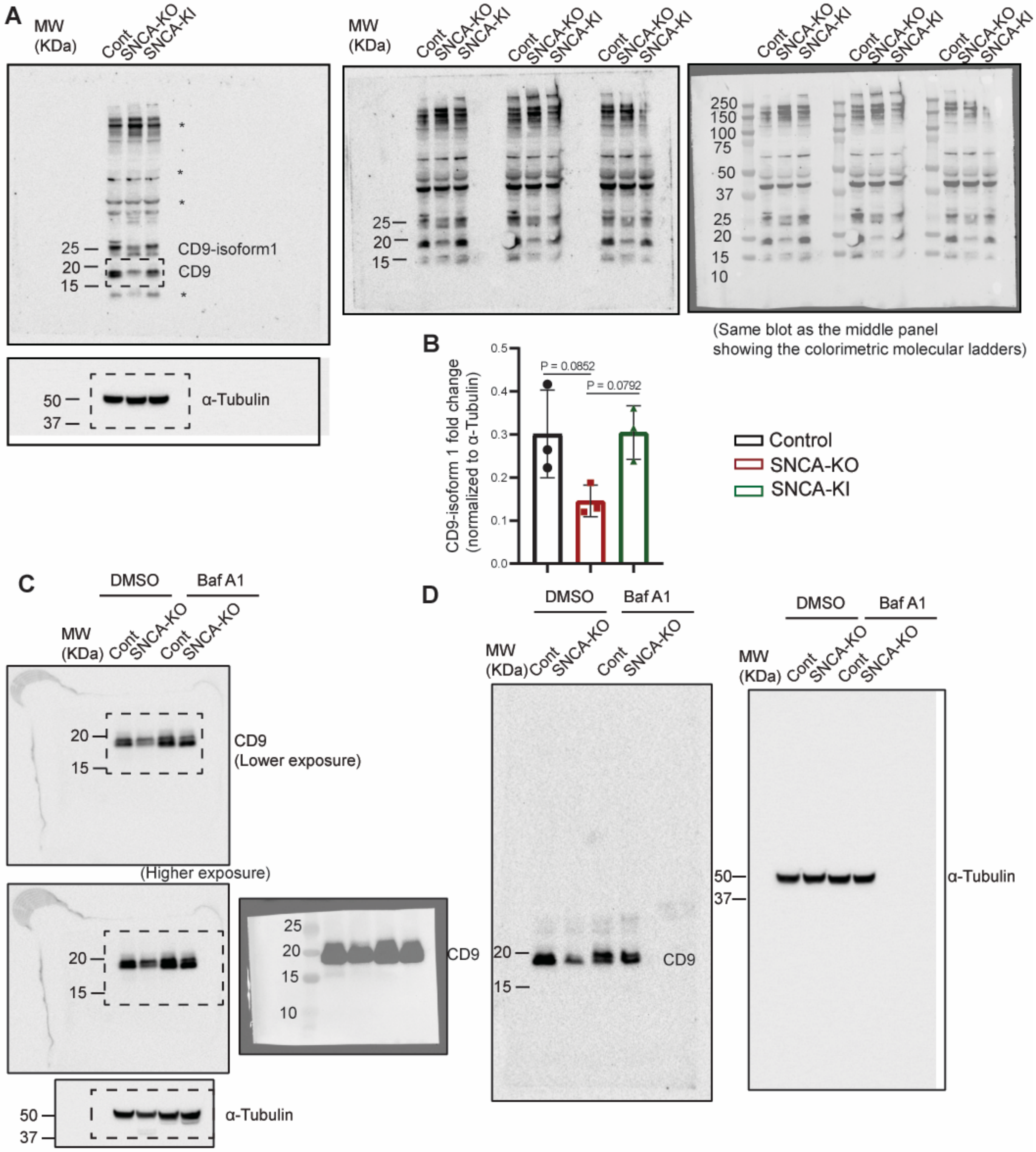
A
MW (KDa)
Cont
SNCA-KO
SNCA-KI
25
20
15
CD9-isoform1
CD9
*
50
37
α-Tubulin
250
150
100
75
50
37
10
(Same blot as the middle panel showing the colorimetric molecular ladders)
B
CD9-isoform 1 fold change (normalized to α-Tubulin)
P = 0.0852
P = 0.0792
0.5
0.4
0.3
0.2
0.1
0.0
Control
SNCA-KO
SNCA-KI
C
DMSO
Baf A1
CD9 (Lower exposure)
(Higher exposure)
D

Figure S5

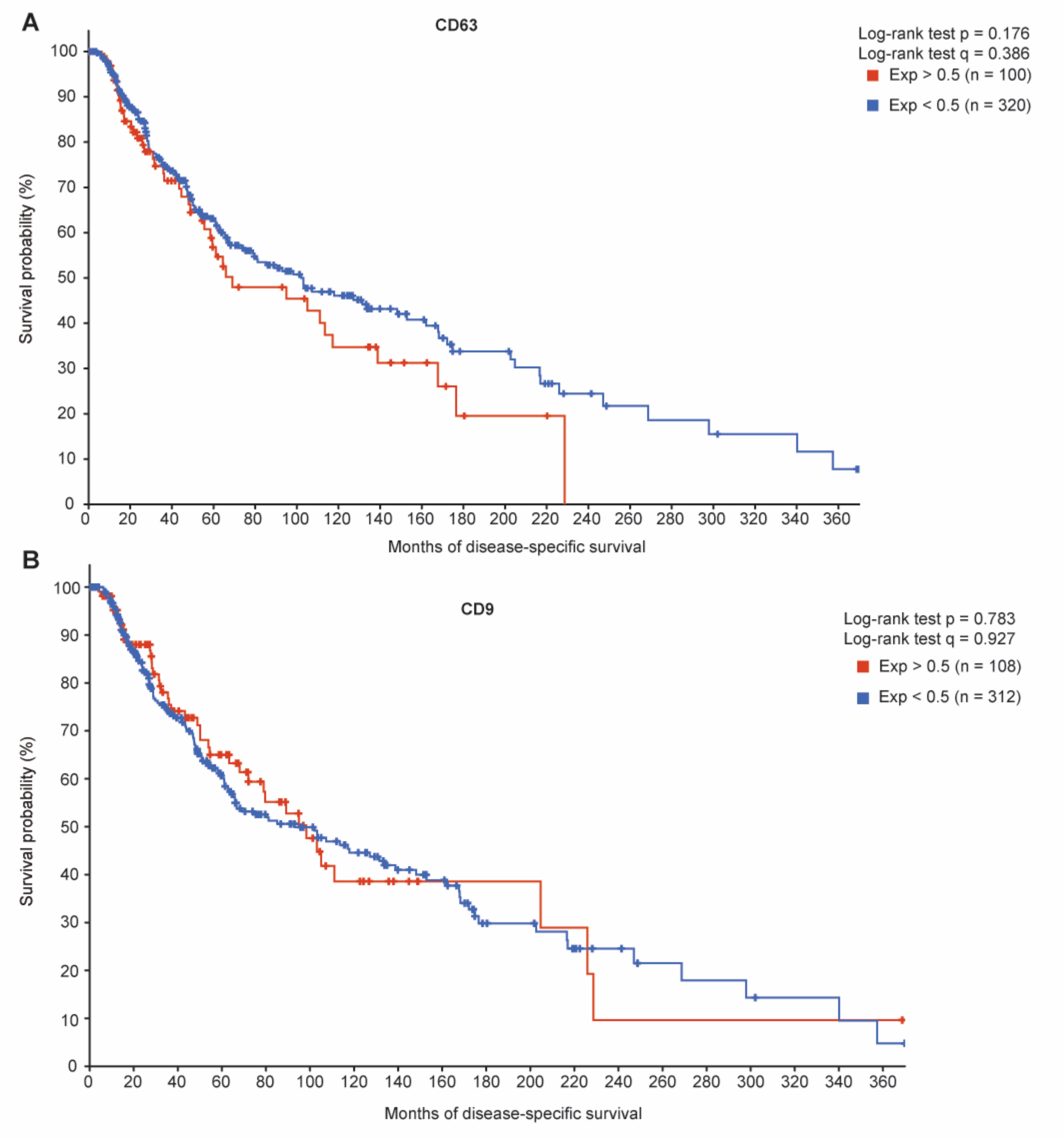
A
CD63
Log-rank test p = 0.176
Log-rank test q = 0.386
Exp > 0.5 (n = 100)
Exp < 0.5 (n = 320)
Survival probability (%)
0
10
20
30
40
50
60
70
80
90
100
0
20
40
60
80
100
120
140
160
180
200
220
240
260
280
300
320
340
360
Months of disease-specific survival
B
CD9
Log-rank test p = 0.783
Log-rank test q = 0.927
Exp > 0.5 (n = 108)
Exp < 0.5 (n = 312)
Survival probability (%)
0
10
20
30
40
50
60
70
80
90
100
0
20
40
60
80
100
120
140
160
180
200
220
240
260
280
300
320
340
360
Months of disease-specific survival
